# Scorpion toxin peptide BMK86-P1 achieves mutation-reversible inhibition of KCNA2 at the cost of reduced efficacy in heteromers and murine neurons

**DOI:** 10.64898/2026.08.28.747831

**Authors:** Elisabeth M.M. Brand, Lorenz Over, Hubert Kalbacher, Remy Lambert, Stefano Iavarone, Hang Lyu, Ulrike B.S. Hedrich, Peter Müller-Wöhrstein

**Author notes:** **Correspondence:** Peter Müller-Wöhrstein.

## Abstract

The discovery of distinctive function-phenotype relationships in monogenetic channelopathies has turned out to be critical for the development of precision medicine approaches. However, the best prediction of clinical phenotypes depends on neuronal function, where existing models lack tools to isolate currents of individual voltage-gated potassium channel subunits and differentiate variant effects in complex systems. Ideally, one should be able to overexpress subunit variants with an additional mutation that confers resistance against the tool to isolate the variant effect. Therefore, we solid-phase synthesized the K_V_1.2 specific scorpion toxin peptide BMK86-P1 and oxidized it with modest efficacy. In mammalian cells this BMK86-P1 selectively inhibited K_V_1.2 homomers, but not heteromers with K_V_1.1. Critically, the *KCNA2* p.Val381Tyr mutation, which reverses BMK86-P1’s selective inhibition of K_V_1.2, also altered the activation of K_V_1.2 homomers to resemble those of K_V_1.1. In addition, BMK86-P1 in murine neurons did not alter passive membrane properties, single action potential properties, or action potential firing. Surprisingly, it induced only minimal changes in spontaneous excitatory postsynaptic currents. In summary, this K_V_1.2 subunit selective toxin peptide asserts its effects primarily on homomeric channels, while only weakly inhibiting K_V_1.2-heteromeric channels and consequently preventing any meaningful impact on neuronal function. This highlights the limits of peptide synthesis together with the need for testing specific compounds on complex systems.

## Introduction

In recent years, advances in genetic diagnostics have highlighted the role of fast-activating voltage-gated potassium channel subunits of the K_V_1 family in neurological disease (Helbig et al., 2016; Paulhus & Glasscock, 2023; Soldovieri et al., 2024; Syrbe et al., 2015). K_V_1 channels encoded by *KCNA1-4* play an important role in brain neurons where they are found in the distal axon initial segment (Inda, DeFelipe, & Muñoz, 2006), the nodes of Ranvier (Rasband & Shrager, 2000) and in presynaptic axon terminals (Trimmer, 2015). In the axon, they are essential for fast repolarization (Kole, Letzkus, & Stuart, 2007) and without them, interneuronal high-frequency action potential firing could not be sustained (McCormick, Connors, Lighthall, & Prince, 1985) and presynaptic excitability would be disturbed (Southan & Robertson, 1998). Structurally, K_V_1 channels are tetramers and assemble as homo- and heteromeric complexes (Long, Tao, Campbell, & MacKinnon, 2007) – in brain neurons typically from K_V_1.1, K_V_1.2 and K_V_1.4 subunits (Manganas & Trimmer, 2000). The inclusion of K_V_1.4 results in a fast-inactivating A-type potassium current, whereas heteromers of K_V_1.1 and K_V_1.2 conduct a slowly inactivating D-type potassium current (Ranjan et al., 2019).

Variants in K_V_1.1 (*KCNA1)* are typically associated with episodic ataxia and epilepsy whereas variants in K_V_1.2 (*KCNA2*) have emerged in developmental and epileptic encephalopathies (DEE). Both loss-of-function or gain-of-function effects have been found either subunit (Masnada et al., 2017; Paulhus & Glasscock, 2023; Syrbe et al., 2015). A pharmacological approach targeting DEE patients with gain-of-function variants of K_V_1.1 and K_V_1.2 has been shown to be successful (Hedrich et al., 2021; Müller et al., 2023) and further approaches are currently being developed (Huang et al., 2024; Snowball et al., 2019), emphasizing the importance of understanding how K_V_1 channels influence neuronal function.

As the influence on neuronal function is very hard to predict by looking at the biophysical properties of a subunit variant alone, variant function should be evaluated directly in a complex system. Ideally, one would want to isolate the effect of K_V_1 variant overexpression in neurons by pharmacological tools that block only wild-type K_V_1 subunits but leave disease-associated variant alone. Then, one could draw clear comparisons with an overexpressed wildtype without having to worry about complex interactions with endogenous expressed potassium channels. Several toxins have been shown to target K_V_1.2 (Harvey & Robertson, 2004; Hopkins, 1998; Sprunger, Stewig, & ÓGrady, 1996). However, no corresponding channel subunit mutation has been described that selectively confers toxin insensitivity to them. Interestingly, the broad potassium channel blocker tetraethylammonium (TEA) gains sensitivity for K_V_1.2 after introduction of the *KCNA2* p.(Val391Tyr) (V381Y) mutation (Al-Sabi et al., 2010). In the last years, this very mutation has been described to reverse the selective action of scorpion toxin peptide mesomatoxin derived from *Mesobuthus martensii* on K_V_1.2 (X. Wang, Umetsu, Gao, Ohki, & Zhu, 2015). The related BMK86 and the even higher K_V_1.2 specific degradation toxin peptide BMK86-P1 share this feature (Mao et al., 2007; Qin et al., 2021; X. Wang et al., 2015). Introducing V381Y alongside human disease-associated variants in an overexpression system may enable selective silencing of endogenous K_V_1.2, allowing functional analysis of K_V_1.2 variants in isolation, as previously implemented with TTX-resistant voltage-gated sodium channels (Lyu et al., 2023).

In this study, we employ the commercially viable Fmoc-/tBu strategy to synthesize the toxin peptide BMK86-P1 and oxidize it due to three disulfate interactions. We were then able to reproduce the inhibitory effect of BMK86-P1 on K_V_1.2 homomers, but found it to be rather ineffective on K_V_1.1/ K_V_1.2 heteromers. In addition, the *KCNA2* V381Y mutation altered the activation kinetics by shifting them closer to K_V_1.1. Finally, BMK86-P1 did not influence action potential properties in primary cortical neurons but preserved spontaneous synaptic activity over time.

## Materials and Methods

### Peptide synthesis

The cyclized peptide ACSKPCRKYCILKKGARNGKCINGRCHCYY was synthesized using the Fmoc-/tBu strategy as described in (Ali et al., 2016; X. Chen, Kalbacher, & Grunder, 2005) with the following modifications. The crude toxin was purified by RP-HPLC on a C18 semi-preparative column (10 × 150 mm; ReproSil 100, Dr. Maisch) using a 40-min gradient of acetonitrile in 0.055% trifluoroacetic acid (0–60% B in 40 min, where B is 80% acetonitrile/H_2_O/0.05% trifluoroacetic acid). This version is referred to as the reduced toxin. Oxidation of the reduced toxin was achieved by dissolving the purified peptide with 2 M acetic acid, and diluted to a peptide concentration of 0.015 mM in the presence of reduced/oxidized glutathione (molar ratio of peptide/GSH/GSSG was 1:100:10) and 2 M guanidine hydrochloride. The solution was adjusted to pH 8.0 with aqueous NH_4_OH and stirred slowly at 4°C for 3 d. The folding reaction was monitored by analytical HPLC. The solution was concentrated using three C18 SepPak (Waters) cartridges connected in series and finally lyophilized. Purification of the oxidized product was achieved first by chromatography on the C18 column using the system above and yielding a purity of ∼80%. Finally, the product was highly purified on a Vydac YMCbasic column using a 60-min gradient, resulting in a purity of ≥95%. The quality of the product was confirmed by analytical HPLC, matrix-assisted laser desorption/ionization time of flight mass spectrometry (MALDI-MS), and electrospray ionization mass spectrometry (ESI-MS), giving the correct mass of oxidized product. The product was diluted in distilled water to a final stock concentration of 1 mM, aliquoted and stored at -20°C.

### *KCNA* plasmids

We employed plasmids with human *KCNA1* or *KCNA2* linked with a P2A linker to RFP or GFP, respectively. For *KCNA1* we used a pcDNA3.1 plasmid purchased from GeneScript (Netherlands) and then ligated with a C-(K)DYK-P2A-tRFP tag by Mahmoud Koko. For *KCNA2* we used a pIRES-AcGFP1 backbone, into which Mahmoud Koko inserted *KCNA2* via overlap PCR. The V381Y mutagenesis was performed using a QuikChange PCR with Pfu polymerase (Promega, Germany).

### Stable *KCNA2* HEK cell line

HEK293 cells (RRID:CVCL_0045) were cultured at 37 °C with 5% CO_2_ in a humidified incubator and grown in Dulbecco’s modified Eagle nutrient medium (Invitrogen, MA, USA) containing 10 % (v/v) fetal calf serum (FCS, PAN-Biothech GmbH, Germany) and 1% L-Glutamine 200 mM (Biochrom GmbH, Germany). They were transduced by Niklas Schwarz with an AAV carrying human *KCNA2* under the CMW promoter with an IRES linked GFP and a puromycin resistance. 6 passages after puromycin selection clones were screened for expression potassium currents by patch clamp and gently frozen in liquid nitrogen for further use. For experiments cells were thawed, passaged at minimum 3 times, split into 3.5 cm wells, rested for 24 h and transferred to the patch-clamp setup.

### Transfection in CHO cells

CHO-K1 cells (RRID:CVCL_0214) were cultured at 37 °C with 5% CO_2_ in a humidified incubator and grown in Ham’s F12 containing 10 % FCS. For transfection, cells were split into 3.5 cm wells and then left to rest for 24 h. 250 µl of Opti-MEM™ (ThermoFischer, Germany) with 2 mg of plasmid and 6 µg polyethylenimine (PEI) as the transfection agent was added to the medium. After 24 h cells were treated with Accutase™ (ThermoFischer, Germany) to detach from the well and were then rested for 30 min to form round attached cells before they were transferred to the patch-clamp setup.

### Primary neuronal cultures

Primary cortical neurons were prepared from mouse embryos of pregnant C57BL/6N mice at embryonic day (E) 18. Experiments were approved by the local Animal Care and Use Committee (Regierungspräsidium Tübingen). Preparation and dissociation of primary cortical neurons was performed as described previously (Hedrich et al., 2021). Cortical neurons were cultured in neurobasal culture medium (Thermo Fisher) with B27^TM^ supplement (Thermo Fisher) and 5 mM L-glutamine (Invitrogen) on microscope coverslips in 24-well cell culture plates (Corning Inc.) at a cell density of 1.5*10^5^ cells/well. To allow adherence of primary cortical neurons, microscope coverslips (VWR International) were precoated with 0.01% poly-D-lysine (Sigma-Aldrich) for 2 h at room temperature. The neuronal growth medium was changed every other day and cells were maintained at 37 °C and 5% CO_2_ in a humidified atmosphere until further experiments. Whole-cell patch clamp recordings on primary pyramidal cortical neurons were performed on *day in vitro* 14 to 17.

### Whole-cell patch clamp

HEK 293, CHO cells and neurons were recorded using an Axopatch 200B amplifier (Molecular Devices), a Digidata 1550B digitizer (Molecular Devices), a micromanipulator SM-5 (Luigs & Neumann), an Axiovert A1 inverse microscope (Zeiss) and pClamp 11.0.3 software (Molecular Devices). Pipettes were pulled from borosilicate glass with a filament with a resistance of 2-3 MΩ for CHO cells and 3-5 MΩ for neurons. HEK 293 and CHO cells were recorded with intracellular solution containing (in mM): 90 KF, 10 KCl, 1 CaCl_2_, 1 MgCl_2_, 10 HEPES, 11 EGTA and 2 Mg-ATP, adjusted to 300 mosmol/l with Mannitol and to a pH of 7.4. For neurons the intracellular solution consisted of the following (in mM): 135 K D-gluconate, 4 NaCl, 0.5 CaCl_2_, 10 HEPES, 5 EGTA, 2 Mg-ATP, 0.4 GTP-Na at 290 mosmol/l and a pH of 7.3. While recording, HEK 293 and CHO cells were maintained in extracellular bath solution containing (in mM): 135 NaCl, 5 KCl, 2 CaCl_2_, 2 MgCl_2_, 5 HEPES and 10 glucose and was adjusted to 320 mosmol/l with a final pH of 7.4. For neurons the extracellular solution consisted of the following (in mM) 140 NaCl, 4.2 KCl, 1 CaCl_2_, 1 MgSO_4_-7H_2_O, 0.5 Na_2_HPO_4_, 0.45 NaH_2_PO_4_, 5 HEPES, 10 glucose, which was adjusted to 305 mosmol/l with a final pH of 7.4. All recordings were performed at room temperature, sampled at 100 kHz and low-pass filtered at 10 kHz.

HEK 293 and CHO cells were selected by patching cells with either GFP, RFP or both giving us the respective homomers and the heteromer. Only cells with series resistance <10 MΩ and a leak <100 pA were selected for recording. Voltage-clamp recordings were performed with whole-cell compensation at 85%. Cells were held at -80 mV in between voltage protocols. Potassium currents were recorded with a 2 s long step protocol which featured +10 mV increments from -80 mV until 60 mV and was followed by a hyperpolarizing step to -120 mV. After the first potassium current protocol 500 µl of Extra were added to CHO cells and the protocol was repeated every 3 mins for a total of 5 recordings. Then, 6 µl of 1 mM BMK86-P1 stock in 500 µl extracellular solution were added to the bath for a final concentration of 3 µM BMK86-P1 and the voltage protocols were run again. The total recording time was 33 mins, during which access resistance was steadily controlled. Extended wash-in periods were tested, but no effect on efficacy was observed (n=3). Reduced or oxidiced BMK86-P1 was added to HEK 293 cells in increasing concentrations ranging from 0.003 µM to 30 µM, following the first potassium current protocol. This protocol was repeated after each addition of the toxin.

Neurons were selected based on their typical pyramidal morphology, a stable resting membrane potential in current clamp mode, a series resistance <20 MΩ and a leak <100 pA. The resting membrane potential was obtained immediately after cell opening. Cells were recorded for at least 2 min clamped at -70 mV to allow the cell to stabilize before starting further protocols. No correction was made for the liquid junction potential. A control baseline recording was first obtained, followed by the application of 3 µM BMK86-P1 or extracellular solution by pipette. After an incubation time of 5 min the neurons were re-recorded.

Passive membrane properties were assessed by current step pulses ranging from -10 pA to - 110 pA, with an increment of -10 pA. Action potential properties were analyzed from the first AP evoked by a 4 s long ramp pulse at a rate of 50 pA/s to accurately determine AP properties. Additionally, a stepwise current injection with an increment of +25 pA and each step lasting 800 ms was applied to determine neuronal firing. To record sEPSCs 50 µM of picrotoxin was added to the extracellular bath solution. Neurons were voltage-clamped at -70 mV and recorded for 2 min with series resistance compensated up to 85%.

### Data analysis and statistics

Data analysis and plotting was performed in pClampFit (v10.7; Molecular Devices 2016) and R (v4.3.3; R Core Team 2024).

Dose-response curves for oxidised and reduced toxin peptide were fitted to the following Hill equation:

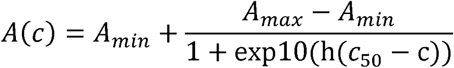

with A being the current peak during BMK86-P1 perfusion divided by the current peak during baseline, *A_min_* and *A_max_* being the minimal and maximal values of this quotient respectively, *c* being the log10 of the concentration of BMK86-P1, *c*_50_ the log10 of the concentration of half-maximal-inhibition and *h* being the Hill slope factor.

The maximum value of the potassium current for extracellular recordings was normalized to the maximum of the baseline recording (Fig. 1C). The maximum of the BMK86-P1 currents were normalized to the maximum of the last extracellular recording (Fig. 1C). Activation kinetics were obtained by first calculating the conductance (g) from the maximum current (I) by dividing it through the difference of the voltage (V) and the equilibrium potential for potassium (V_E_ = -75 mV).

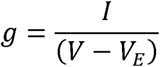

**Figure 1.**
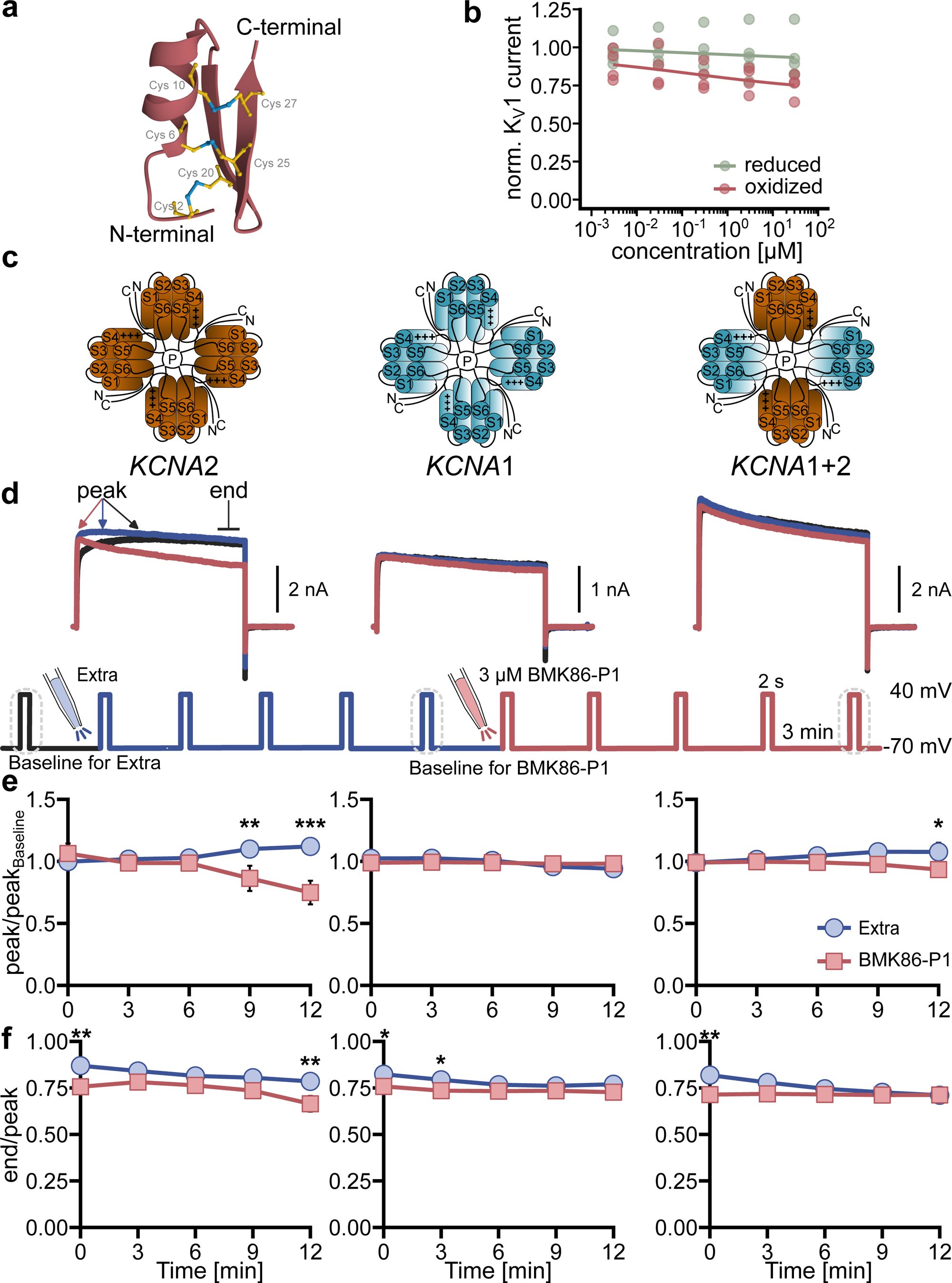
Effects of BMK86-P1 on K_V_1.2 and K_V_1.1 tetramers (A) Three-dimensional structure of the oxidized BMK86-P1 modeled with AlphaFold3. Disulfide bridges (violet) between cysteine residues (yellow) are shown. (**B**) Dose-response curve for reduced or oxidized BMK86-P1 at increasing concentrations of 0.003 µM, 0.03 µM, 0.3 µM, 3 µM, 30 µM recorded in HEK 293 cells stably transfected with human K_V_1.2. Normalized potassium currents were obtained by dividing the peak current at the respective concentration by the baseline. (**C**) Sketches of the K_V_1 channel with different compositions. Each subunit consists of 6 transmembrane domains, where the S4 domain carries the voltage sensor (indicated by +++) and the linkage between S5 and S6 forms the pore (P). (**D**) Top: Example potassium currents recorded in CHO cells transfected with *KCNA2* (left), *KCNA1* (middle) or both (right). ‘peak’ and ‘end’ highlight the peak of the current and the steady-state current at the end of the voltage step respectively. Bottom: Voltage protocol for recordings. Dotted circles mark steps of the examples above. (**EF**) Normalized potassium current obtained by dividing peak by the respective baseline (**E**) or inactivation as measured by dividing end by peak (**F**) after the application of extracellular solution (blue) and 3 µM BMK86-P1 (red) for different compositions.

Afterwards conductances were normalized and fit to

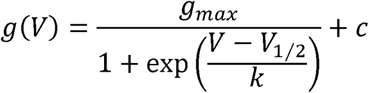

with g_max_ being maximal normalized conductance, V voltage, V_1/2_ voltage at half activation, k slope factor and c a constant term. The inactivation ratio was obtained by dividing the mean steady state current of the last 50 ms of the voltage step through peak current.

For AP firing frequency, events exceeding the 0 mV threshold were detected in the stepwise current-injection protocol. AP properties were obtained from the first derivative of the first evoked AP of the ramp recording. The voltage threshold and rheobase were determined as the voltage and injected current when the derivative crossed over 10mV/ms. Amplitude and rising time were calculated from the voltage threshold to the action potential peak. FWHA was defined as the time at which the AP was more positive than the half maximum amplitude. In addition, the repolarizing time was derived from the voltage peak to the minimum potential and the AHP from the difference between the minimum potential and the baseline voltage. For the evaluation of sEPSC files were digitally low-pass filtered at 1000 Hz and identified using custom spike detection scripts in Spike 2 (v10.22, Cambridge Electronic Design 2024).

For statistical analysis, all data were tested for normality using the Shapiro-Wilk test with a significance level of 0.05. All data of HEK 293 and CHO cells are given as mean ± SEM. We employed two-way ANOVA with appropriate post-hoc tests for the comparisons in CHO cells. To evaluate the effect of BMK86-P1 on primary cortical cultures, Wilcoxon-signed rank test was used for non-normally distributed data. A mixed effect model with linear regression was employed for the cumulative plots of sEPSC amplitude and instantaneous frequency with random effects assigned for each cell. The data are given as marginal means ± SE together with the t-value. All other neuron data are given as median and IQR = Q1 – Q3. Significance with respect to control is indicated in the figures as follows: *p < 0.05, **p < 0.01, and ***p < 0.001.

## Results

### The selectivity of BMK86-P1 is exerted on *KCNA2* homomers

BMK86-P1 was synthesized using the Fmoc-/tBu strategy, purified by RP-HPLC, oxidized, purified again and finally verified using analytical HPLC, matrix-assisted laser desorption/ionization time of flight mass spectrometry (MALDI-MS), and electrospray ionization mass spectrometry (ESI-MS) (see Methods). The tertiary structure of BMK86-P1 features three disulfide bonds (Fig 1A) which likely have a key function in stabilizing its structure. To confirm the formation of these disulfide bonds, we first determined the biological activity of the peptide by comparing the reduced and oxidized version in HEK 293 cells stably transduced with *KCNA2*. Here we saw that indeed the reduced toxin peptide hardly saw any effect on potassium current while we observed a concentration dependent reduction of ∼25% for the oxidized version (Extra sum-of-squares F-test, F = 5.8, p < 0.001, n = 4 or 5, see Table 1, Fig. 1A-B).

**Table 1.**
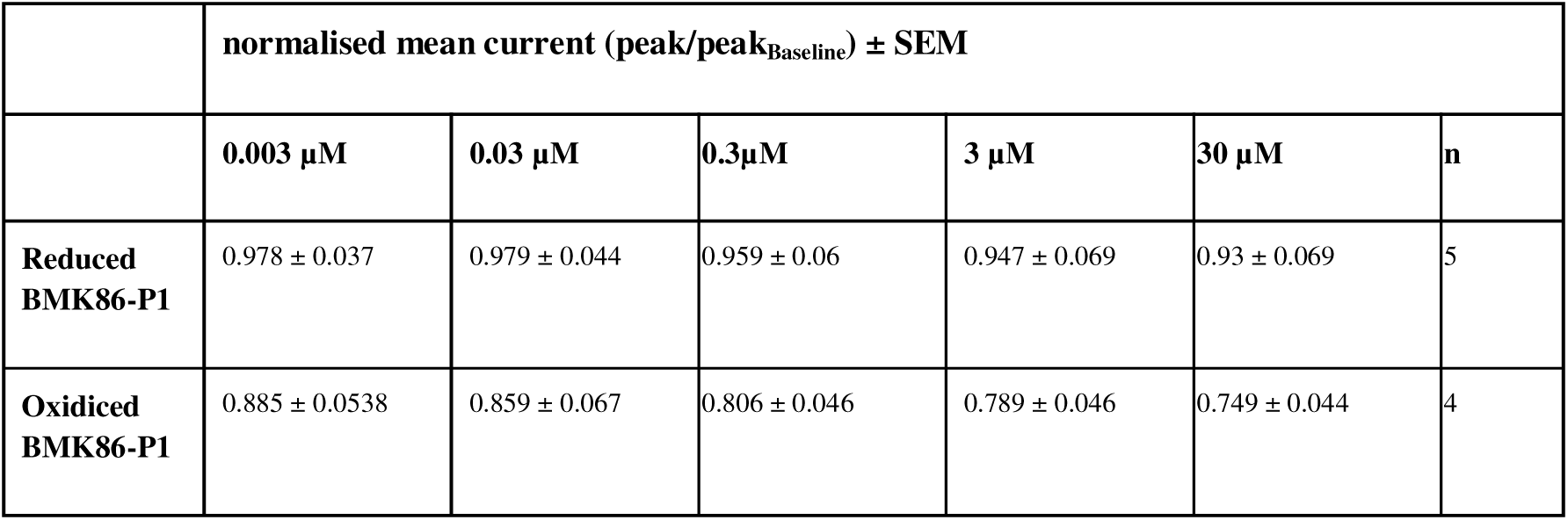
Reduced or oxidiced BMK86-P1 effect on *KCNA2*. Summary for the last recorded step after addition of increasing concentrations of the respective toxin. Potassium peak currents were normalized to baseline peaks.

As we reasoned that the mild effect of the oxidised toxin peptide (compared to an IC_50_ of 29 nM (Qin et al., 2021)) might partially be caused by endogenous expression of other toxin insensitive potassium currents in HEK cells, for the following experiments CHO cells were chosen which lack endogenous potassium currents (Hebert, Desir, Giebisch, & Wang, 2005). To assess the selectivity of the oxidized BMK86-P1 on homo- and heteromers of K_V_1.2, we employed transient transfections with DNA of either *KCNA1*, *KCNA2* or both (Fig. 1C). Cells were recorded for a total of 12 min after peptide application and inhibition was assessed by step pulses to +40 mV every 3 min. To exclude potential rundown and/or use-dependence effects (Rezazadeh, Kurata, Claydon, Kehl, & Fedida, 2007), all cells were first recorded after the addition of extracellular fluid (Extra), followed by a repeat of the recordings under application of the peptide (see Fig. 1D). Control experiments with two consecutive applications of Extra showed no differences in the potassium currents of K_V_1.2 channels over time course of the recording (1.277 ± 0.157 vs 0.960 ± 0.128 for Extra and BMK86-P1 respectively, p=0.131, Two-way ANOVA with Tukey Post-hoc test, n=5, Fig. S1A).

3 µM BMK86-P1 reduced the homomeric K_V_1.2 mediated potassium current by 33 % 12 min after application (see Table 2, Fig. 1E). To examine potential effects of the peptide on inactivation we calculated a proxy for inactivation by dividing the current at the end of the voltage step by its peak. This was necessary as not all recordings saw sufficient inactivation occur within the voltage step to fit an exponential decay. In the end/peak ratio values closer to 1 suggest missing inactivation either due to a late peak current or due to no inactivation occurring, while values closer to 0 imply full inactivation. Here we saw a 50% relative increase of the portion of inactivated current for K_V_1.2 after 12 min (see Table 2, Fig. 1F). In homomeric K_V_1.1 channels, no significant change in current amplitude was detected after 12 min (see Table 2, Fig. 1E) and inactivation was not significantly changed after 12 min (see Table 2, Fig. 1F). The difference at 0 min is most likely caused by a time depend rundown effect as the ratios are very similar for 12 min of Extra (0.770 ± 0.018) and 0 min of BMK86-P1 (0.758 ± 0.024). Finally, we performed co-transduction of *KCNA1* and *KCNA*2 and found a decrease of 13% in potassium current (see Table 2, Fig. 1E). With regards to the inactivation ratio, no significant difference was detected after 12 minutes (see Table 2, Fig. 1F), while a similar rundown effect to *KCNA1* can be observed overall.

**Table 2.** BMK86-P1 effect on *KCNA2*, *KCNA2* V381Y, *KCNA1* and *KCNA1*+*KCNA2*. Summary for the last recorded step at 12 min. Potassium peak currents were normalized to respective baseline peaks. A proxy for inactivation was obtained by dividing the steady state current at the end of the voltage step by the peak current. p-values were obtained using a Two-way ANOVA with Tukey post-hoc testing.

|  | normalised current (peak/peak<sub>Baseline</sub>) |  |  | inactivation ratio (end/peak) |  |  | n |
| --- | --- | --- | --- | --- | --- | --- | --- |
|  | Extra | BMK86-P1 | p | Extra | BMK86-P1 | p |  |
| **KCNA2** | 1.120 ± 0.053 | 0.748 ± 0.095 | **p<.0001** | 0.785 ± 0.026 | 0.664 ± 0.042 | **p=0.003** | 15 |
| **KCNA1** | 0.941 ± 0.045 | 0.981 ± 0.027 | p=0.358 | 0.770 ± 0.018 | 0.727 ± 0.058 | p=0.121 | 11 |
| **KCNA1 + KCNA2** | 1.076 ± 0.055 | 0.934 ± 0.028 | **p=0.015** | 0.711 ± 0.027 | 0.712 ± 0.013 | p=0.968 | 7 |
| **KCNA2 V381Y** | 0.761 ± 0.065 | 0.743 ± 0.066 | p=0.884 | 0.885 ± 0.031 | 0.790 ± 0.059 | p=0.209 | 7 |

Given that K_V_1 channel subunits assemble as tetramers, we calculated the expected effects of different tetrameric compositions of K_V_1.1 and K_V_1.2 as they would appear by Poisson’s law (Fig. S1A) under various assumptions of subunit-inhibition interaction. From these models, it is unlikely that either only K_V_1.2 homomers are inhibited or inclusion of only one subunit is sufficient for a full block (Fig. S1B).

Taken together, these findings indicate that BMK86-P1 is a specific blocker of K_V_1.2 homomers and loses efficacy in heteromers.

### The toxin resistance mutation V381Y impacts K_V_1.2 gating

The toxin resistance conferring variant is located in the pore region of *KCNA2* and substitutes the valine residue in *KCNA2* with the tyrosine found at the paralogous position 379 in *KCNA1* (Fig. 2A). We could replicate the reduced efficacy of BMK86-P1 on *KCNA2*-V381Y (see Table 2, Fig. 2B). However, it remains unclear how this mutation impacts the kinetics of K_V_1.2. The in-silico tool ‘perfeKt’ predicted a gain-of-function for this variant (Bosselmann et al., 2022), which was confirmed experimentally by a ∼8 mV leftward shift of the half-activation voltage(V_1/2_) (see Table 3, Fig. 2C and D).

**Figure 2.**
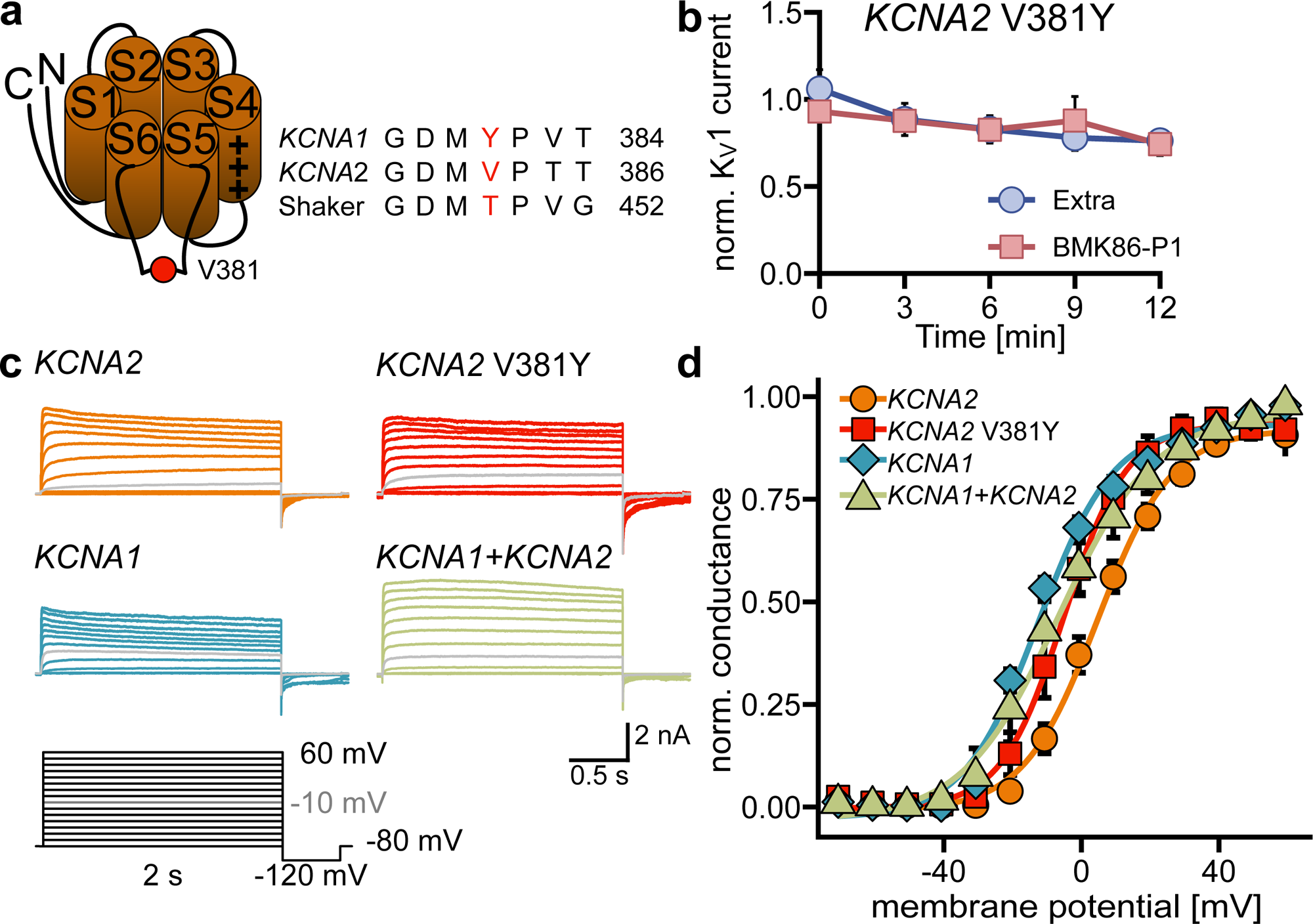
Impact of the V381Y mutant on K_V_1.2 activation (**A**) Schematic representation of the K_V_1.2 subunit with highlight (red) on the Valine at position 381 and comparison to paralogous sequences. (**B**) Normalized potassium currents at 40 mV recorded after application of extracellular solution and 3 µM BMK86-P1. (**C**) Representative current traces of activation protocols recorded from cells transfected with *KCNA2*, *KCNA1*, *KCNA1+KCNA2* or the *KCNA2* V381Y mutant (left). The pulse to -10 mV is marked in grey. (**D**) Activation curves of cells transfected with *KCNA2*, *KCNA2* V381Y, *KCNA1* and *KCNA1*+*KCNA2*.

**Table 3.** Activation characteristic for *KCNA2*, *KCNA2* V381Y, *KCNA1* and *KCNA1*+*KCNA2*. Values were obtained as means of individual fits of normalized conductance to a Boltzmann equation with g_max_ being maximal normalized conductance, V_1/2_ voltage at half activation, k slope factor and c as a constant term (see Methods). p-values for pairwise t-tests of V_1/2_, due to multiple testing the threshold for significance (marked in bold) was Bonferroni adjusted to an α of 0.017.

| | $V_{1/2}$ | $k$ | $g_{\max}$ | $c$ | $p(V_{1/2})$ vs *KCNA2* | n |
| --- | --- | --- | --- | --- | --- | --- |
| *KCNA2* | $6.80 \pm 2.12$ mV | $10.79 \pm 0.75$ | $0.97 \pm 0.02$ | $-0.01 \pm 0.00$ | - | 19 |
| *KCNA2 V381Y* | $-2.73 \pm 3.34$ mV | $8.81 \pm 1.31$ | $0.98 \pm 0.02$ | $-0.01 \pm 0.00$ | **0.012** | 11 |
| *KCNA1* | $-9.93 \pm 1.93$ mV | $12.10 \pm 1.23$ | $1.00 \pm 0.01$ | $-0.02 \pm 0.01$ | **<0.001** | 16 |
| *KCNA1 + KCNA2* | $-5.54 \pm 4.73$ mV | $11.94 \pm 1.06$ | $0.99 \pm 0.01$ | $-0.01 \pm 0.00$ | **0.006** | 7 |

### BMK86-P1 has no detectable effect on neuronal function

Since the toxin is a specific blocker of K_V_1.2 homomers, but only had a mild inhibitory effect on heteromers, we hypothesized that BMK86-P1 might also only slightly alter neuronal function. K_V_1.2 subunits are predominantly found as heteromers in neurons, where they assemble together with other K_V_1 channel subunits (Trimmer, 2015). To test this, we employed primary cortical cultures and applied 3 µM BMK86-P1 of after recording baseline neuronal activity. To exclude potential effects of repeated measurements, a control experiment was performed with the addition of Extra alone (Fig. 3A).

**Figure 3.**
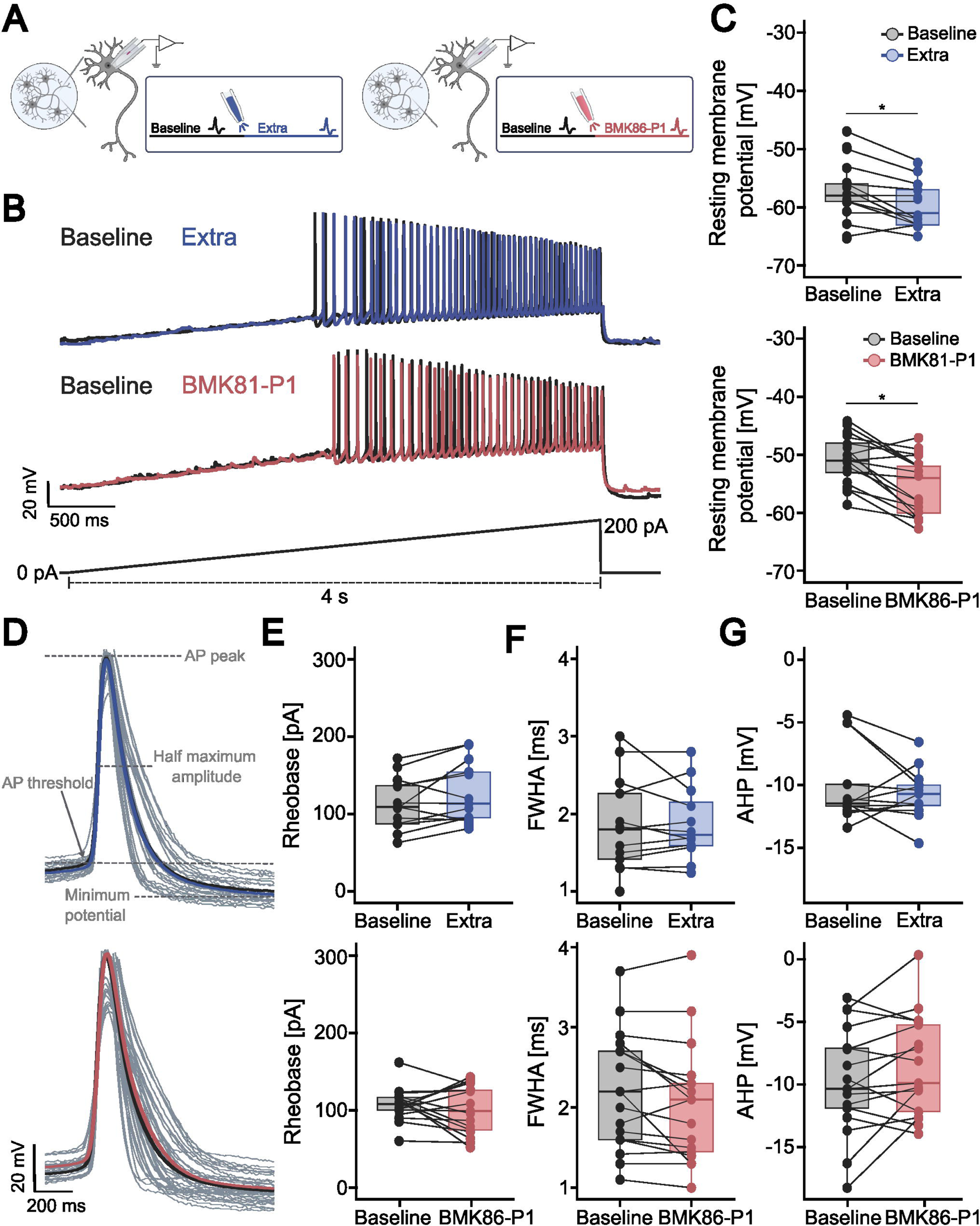
Effect of BMK86-P1 on AP properties of pyramidal cortical neurons. (**A**) Schematic of recording protocol of pyramidal cortical neurons before (baseline, black) and after application of extracellular solution, abbreviated as Extra (blue) or BMK86-P1 (red). After administration of each substance, recording protocols were repeated. (**B**) Representative AP traces of pyramidal cortical neurons treated with Extra (blue) or BMK86-P1 (red) compared to baseline (black) in response to a 4 s ramp pulse at a speed of 50 pA/s. (**C**) Resting membrane potential before (baseline, black) and after application of Extra (blue) or BMK86-P1 (red). (**D**) Average AP waveform of the first evoked AP after a 4 s long ramp pulse at a rate of 50 pA/s before (baseline, black) and after application of Extra (blue) or BMK86-P1 (red). Bold lines represent the average traces, thin gray lines represent the single APs. Landmarks used to calculate the AP parameters are indicated: AP threshold, AP peak, maximum half width and minimum threshold. (**E**) Rheobase before (baseline, black) and after application of Extra (blue) or BMK86-P1 (red). (**F**) FWHA recorded after application of Extra (blue) or BMK86-P1 (red) in comparison with the respective baseline (black). (**G**) AHP amplitude recorded after application of Extra (blue) or BMK86-P1 (red) in comparison with the respective baseline (black).

We found that both BMK86-P1 and Extra application decreased the resting membrane potential by 3 mV (see Table 4, Fig. 3C), indicating a toxin-independent effect. Passive membrane parameters including input resistance and rheobase were unaltered between Extra or BMK86-P1 recorded neurons and baseline (see Table 4, Fig. 3E, S2F). In contrast, the action potential (AP) threshold was hyperpolarized by 3 mV after application of both Extra or BMK86-P1 compared to the respective baseline, indicating that the reduction in the AP threshold was time-dependent (see Table 4, Fig. S2E). Further analysis of the AP waveform of the first evoked AP in a ramp recording revealed mostly no changes, as rising time, full width at half amplitude (FWHA), amplitude and repolarizing time were unaltered between baseline and Extra or BMK86-P1 application (see Table 4, Fig. 3DF, S2BCD). Additionally, the afterhyperpolarization (AHP) amplitude was unaffected between Extra or BMK86-P1 and respective baseline (see Table 4, Fig. 3G). Together, these data indicate that BMK86-P1 does not affect intrinsic neuronal excitability under the tested conditions.

**Table 4.** Single AP parameters for primary cortical neurons treated with extracellular fluid or BMK86-P1. Data are presented as median with interquartile range for extracellular fluid recordings and BMK86-P1 recordings, as well as the respective baseline recordings. Wilcoxon signed-rank test was used to evaluate the differences between paired recordings of baseline and extracellular fluid or BMK86-P1.

|  | Control recording |  |  |  |  | BMK86-P1 recording |  |  |  |  |  |  |
| --- | --- | --- | --- | --- | --- | --- | --- | --- | --- | --- | --- | --- |
|  | Baseline |  | Extracellular fluid |  | n | p-value | Baseline |  | BMK86-P1 |  | n | p-value |
| **Resting membrane potential (mV)** | -58 | -59 to -56 | -61 | -63 to -57 | 14 | 0.012 | -51 | -53 to -48 | -54 | -60 to -52 | 17 | 0.001 |
| **Input Resistance (MΩ)** | 171.89 | 137.68 to 257.18 | 170.52 | 108.46 to 199.21 | 14 | 0.092 | 266.39 | 221.25 to 291.28 | 275.3615 | 243.3 to 318.95 | 17 | 0.224 |
| **Rheobase (pA)** | 114.37 | 87.66 to 136.63 | 113.63 | 95.52 to 154.33 | 13 | 0.08 | 108.1 | 100.56 to 116.73 | 99.16 | 74.72 to 126.02 | 17 | 0.487 |
| **AP threshold (mV)** | -39.27 | -42.11 to -36.92 | -42.11 | -43 to -38.45 | 13 | 0.01 | -38.15 | -39.37 to -37.53 | -40.8 | -43.33 to -38.45 | 17 | < 0.001 |
| **Amplitude (mV)** | 88.8 | 83 to 90.33 | 87.28 | 84.22 to 93.38 | 13 | 1 | 91.86 | 70.5 to 100.1 | 93 | 75.38 to 101.32 | 17 | 0.602 |
| **Rising time (ms)** | 0.84 | 0.75 to 0.9 | 0.8 | 0.77 to 0.84 | 13 | 0.944 | 0.94 | 0.84 to 1.1 | 1 | 1 to 1 | 17 | 0.752 |
| **FWHA (ms)** | 1.8 | 1.42 to 2.26 | 1.69 | 1.57 to 2.1 | 13 | 0.968 | 2.2 | 1.6 to 2.7 | 2.1 | 1.45 to 2.3 | 17 | 0.069 |
| **Repolarization time (ms)** | 15.6 | 13.65 to 21.6 | 14.28 | 11.49 to 17.3 | 13 | 0.375 | 17.1 | 11.9 to 20.7 | 16.2 | 12.9 to 25.4 | 17 | 0.587 |
| **AHP (mV)** | -11.47 | -11.67 to -9.95 | -10.71 | -11.61 to -10 | 13 | 0.541 | -10.34 | -11.87 to -7.11 | -9.88 | -12.41 to -5.25 | 17 | 0.159 |

Next, we assessed the AP firing rate, with the firing frequency of Extra-treated neurons being largely unaffected compared to baseline, as shown by the similar area under the curve (2900 [2600 to 4463] vs 2850 [1688 to 3963] for Extra, p = 0.091 Wilcoxon signed-rank test, n = 13 and 275.5 [225.1 to 425.1] vs 307.4 [206.4 to 456.2] for BMK86-P1, p = 0.922, Wilcoxon signed-rank test, n = 17, Fig. 4A). However, BMK86-P1-treated neurons displayed a slight trend towards higher AP frequencies at relatively low current injections, while exhibiting a more pronounced depolarization block at higher current injections.

**Figure 4.**
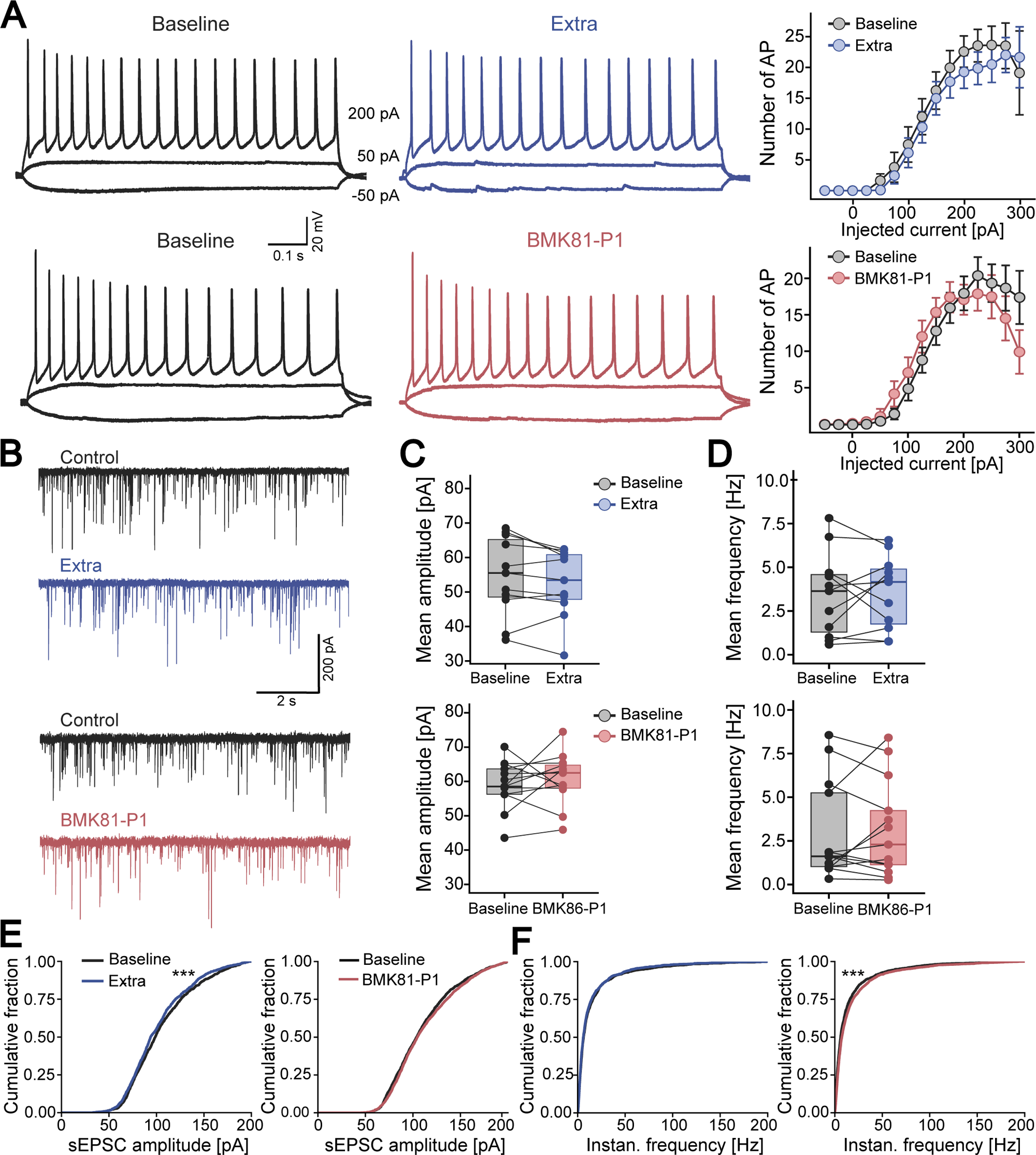
Impact of BMK86-P1 on the excitability of pyramidal cortical neurons and neural network activity. (**A**) Representative traces of AP trains of pyramidal cortical neurons in response to current injections of -50pA, 50 pA and 200 pA pulses for 800 ms before (baseline, black) and after application of Extra (blue) or BMK86-P1 (red). (**B**) Representative traces of sEPSCs recorded from cortical pyramidal neurons before (baseline, black) and after application of Extra (blue) or BMK86-P1 (red). (**C**) Mean sEPSC amplitude recorded from pyramidal cortical neurons before (baseline, black) and after application of Extra (blue) or BMK86-P1 (red). (**D**) Mean sEPSC frequency recorded from pyramidal cortical neurons before (baseline, black) and after application of Extra (blue) or BMK86-P1 (red). (**E**) Cumulative fraction of sEPSC amplitudes recorded over a 2 min period following application of Extra (blue) or BMK86-P1 (red) was compared to baseline (black). (**F**) Cumulative fraction of sEPSC instantaneous frequencies recorded over a 2 min period following application of Extra (blue) or BMK86-P1 (red) was compared to baseline (black).

Finally, we investigated whether small differences in AP frequency impact the activity of neural networks. To this end, we characterized spontaneous excitatory postsynaptic currents (sEPSC) in pyramidal neurons. Neither the mean amplitude nor frequency of sEPSC differed significantly between baseline and either treatment condition (see Table 5, Fig. 4BCD).

**Table 5.**
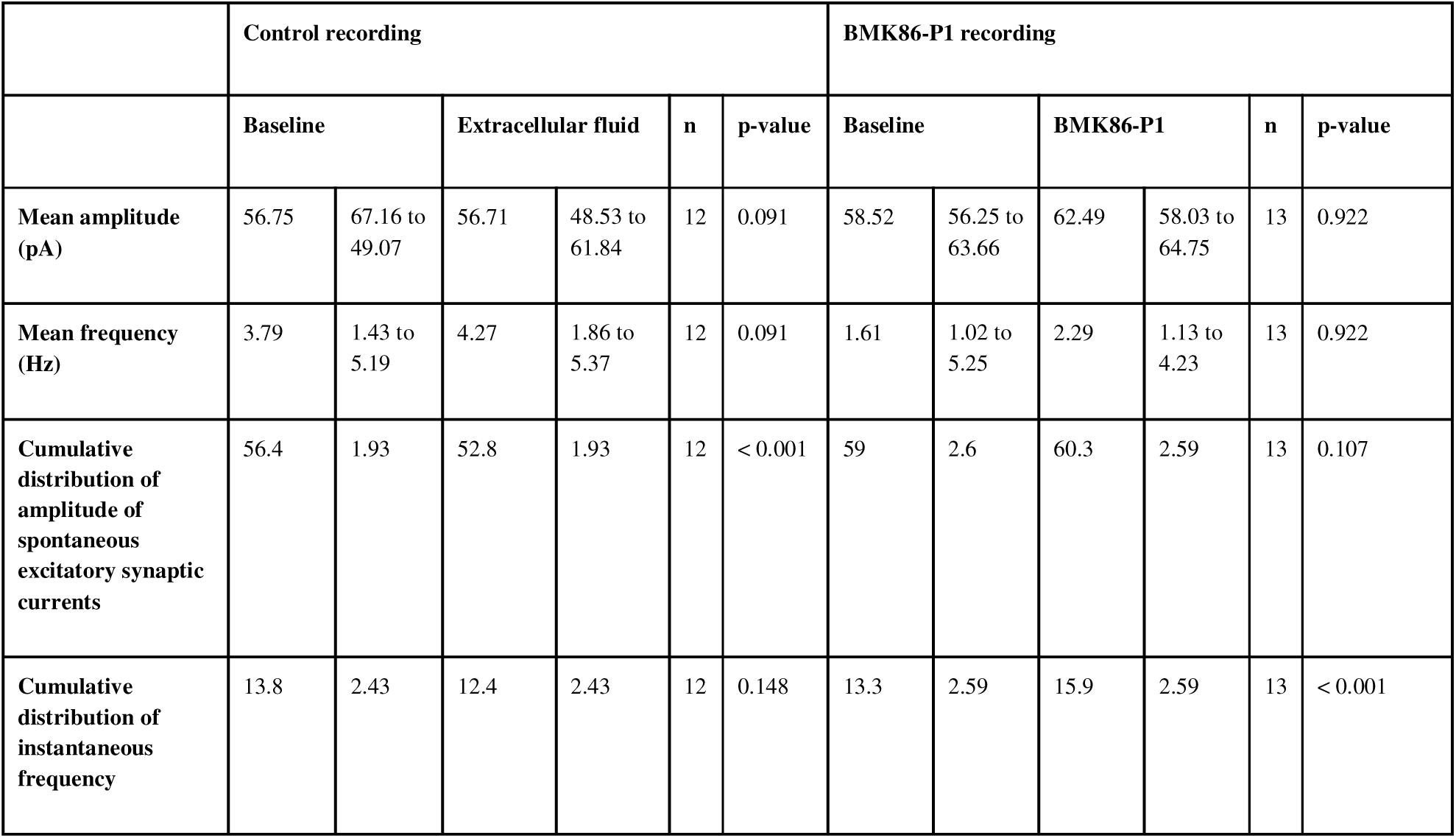
Characteristics of spontaneous synaptic activity for primary cortical neurons treated with extracellular solution or BMK86-P1. Data were obtained as median with interquartile range for extracellular solution recordings and BMK86-P1 recordings, as well as the respective baseline recordings. Values were tested using Wilcoxon signed-rank test to compare paired measurements of baseline and Extra or BMK86-P1. Marginal means ± SE are given for the cumulative distributions of amplitude and instantaneous frequency of spontaneous excitatory synaptic currents. A mixed effect model with linear regression was used to evaluate BMK86-P1-related differences in amplitude and instantaneous frequency, with cell as a random effect.

Similarly, the cumulative distribution of the sEPSC amplitude of neurons treated with BMK86-P1 showed no significant difference compared to baseline, while the sEPSC amplitude of Extra-treated neurons was significantly reduced (see Table 5, Fig. 4E). Interestingly, the instantaneous frequency was unaffected after the application of Extra compared to baseline, while BMK86-P1 resulted in a significant increase in the instantaneous sEPSC frequency. However, this increase was relatively small in comparison to baseline (see Table 5, Fig. 4F).

Altogether, these data indicate no meaningful effect of BMK86-P1 on neuronal excitability and synaptic activity, which is consistent with the selective, but mild blockade of K_V_1.2 homomers, but not K_V_1.2-containing heteromers.

## Discussion

Toxins purified from venomous animals are a rich source for a diverse pool of K^+^ channel blockers. K_V_1 channel blockers have emerged as powerful research tools to localize channel expression pattern, identify their function in physiological systems or characterize structural and biophysical properties of channel pores (Hidalgo & MacKinnon, 1995; Kole et al., 2007; Southan & Robertson, 1998; F. C. Wang, Parcej, & Dolly, 1999). The increasing use of transcriptional and proteomic analysis has led to the discovery of a large number of toxin peptides, which are often highly homologous, but with different pharmacological profiles for specific channel subtypes (Grissmer et al., 1994; Mendes, Viana, Nencioni, Pimenta, & Beraldo-Neto, 2023). In recent years, highly selective Kv1.2 subunit blockers have been described with an increased affinity for K_V_1.2-containing channels than for other K_V_1 channel subunits. (P. Chen, Dendorfer, Finol-Urdaneta, Terlau, & Olivera, 2010; Luna-Ramirez et al., 2014; Rodrigues, Arantes, Monje, Stuhmer, & Varanda, 2003; Shakeel et al., 2025; Visan, Fajloun, Sabatier, & Grissmer, 2004). Most studies have focused on overexpressed K_V_1.2 homomers in heterologous expression systems, while toxin sensitivity in heteromers is expected to depend on subunit stoichiometry and spatial arrangement (Al-Sabi et al., 2010). As K_V_1 channels assemble predominantly in heteromers in physiological conditions, studying toxin-channel formation in physiologically relevant K_V_1 heteromers is essential to predict toxin selectivity (Cordeiro et al., 2019) and will help to identify the physiological role of K_V_1 assemblies. Using complex preparations thus allows assessment of all possible currents that might arise from different compositions of K_V_1 channels, be it A or D type. Moreover, studying the underlying binding mechanisms will help to discover the structural features that regulate toxin sensitivity, as demonstrated with K_V_1.2 and maurotoxin, and could facilitate the development of highly potent and subtype-specific blockers (Visan et al., 2004; Wu et al., 2024). To date, no critical binding site in K_V_1.2 has been identified that would reverse the toxińs affinity.

Here, we investigated the pharmacological properties of the toxin-derived peptide BmK86-P1 by synthesizing it with the Fmoc/tBu strategy (Mao et al., 2007; Qin et al., 2021). Our findings show that BMK86-P1 selectively blocks voltage-gated K_V_1.2 subunits in their homomeric form without affecting the activation or inactivation kinetics. As expected, BMK86-P1 showed no significant inhibitory effect on Kv1.1 homomeric channels. Although the selectivity for K_V_1.2 was preserved, the absolute potency of Kv1.2 inhibition in CHO cells was lower than previously reported, potentially reflecting expression system–dependent differences (Qin et al., 2021) or a failure of disulfide bond formation due to incomplete oxidation. Disulfide brigdes between cysteine residues formed through oxidative processes play a critical role in the correct folding of peptides. These disulfide bonds stabilize the three-dimensional structure, thereby enable the toxińs biological activity (Bulaj, 2005; Gehrmann, Alewood, & Craik, 1998; Li, Schoneich, & Borchardt, 1995). The peptidés native three-dimensional structure is essential for channel-toxin interactions, as proper folding ensures the correct positioning of key binding residues (Pennington et al., 1999; Tabassum et al., 2017). Although the crucial disulfide bridges were introduced during the oxidation process and their formation was verified by analytical HPLC, incomplete oxidation or exposure to reducing environment may have compromised the toxińs high affinity to the channel (Armishaw et al., 2006).

We further demonstrated that the toxin peptide loses its strong effect when K_V_1.2 is co-assembled with K_V_1.1 in heteromeric channels. Computational modelling of different subunit compositions supports these finding that inhibition is not only determined by K_V_1.2 homomers, but suggests that the presence of only Kv1.2 subunits alone is insufficient to achieve full inhibition within tetrameric assemblies. Nonetheless, our findings require further validation by using systems that allow the controlled expression of the K_V_1.2 subunit spatial arrangements and composition within the channel (Al-Sabi et al., 2010; Heginbotham & MacKinnon, 1992).

The introduction of the V381Y mutation in *KCNA2* then leads towards inhibition of K_V_1.2 current with a similar IC_50_ to K_V_1.1 (Al-Sabi et al., 2010). In the case of TEA it is assumed that the aromatic residue of tyrosine in p.Y379 attracts the positively charged nitrogen and thus holds it on top of the entry into the pore blocking potassium ions from entry (Heginbotham & MacKinnon, 1992). The reverse mechanism could underlie the reduced blocking efficacy of BMK86-P1 in opposing K_V_1.2/K_V_1.1 heteromers, although in this case it might rather be a space restriction. Using alphafold3 (Abramson et al., 2024) we found the distance of opposing p.381 in K_V_1.2 tetramers increased by ∼1 Å for valine in comparison to tyrosine (Fig. S3AB), thereby widening the region above the pore. This may allow for sulfur mediated van-der-Waals interactions between p.M380 and the 6 cysteins in BMK86-P1 (Gómez-Tamayo et al., 2016). Opposing p.Y379 in K_V_1.1 tetramer models show a similar distance to *KCNA2* V381Y (Fig. S3C) and the opposing heteromer mimics this as well (Fig. S3D). Consequently, we were able to replicate the selectivity of BMK86-P1 to p.V381 in *KCNA2*. We then demonstrated that the p.V381Y variant shifts the activation towards more hyperpolarized potentials, thereby resembling the activation kinetics of K_V_1.1. This suggests that the amino acid in the filter region of the pore is crucial for the affinity of the toxin, but also contributes to the subunit-specific gating properties of highly homologous K_V_1 channels, as recently emphasized by the closed state Shaker Cryo-EM (Liu, Bassetto, Contreras, Perozo, & Bezanilla, 2025).

Dysfunction of K_V_1.2 channel has been linked to severe neurological disorders (Fulton et al., 2011; Helbig et al., 2016). Reduced K_V_1.2 function, often caused by genetic variants, is particularly associated with epilepsy, intellectual disability and episodic ataxia (Helbig et al., 2016; Masnada et al., 2017; Syrbe et al., 2015). Given the key role of K_V_1.2 in maintaining neuronal excitability (Kole et al., 2007), a reduction in K_V_1.2 activity is expected to promote neuronal hyperexcitability and firing rates, potentially contributing to epileptiform activity. Homozygous K_V_1.2 knock-out mice exhibit early lethality and spontaneous seizures, although many functional features of neurons surprisingly remain unaffected (Brew et al., 2007). This might be due to rearrangement of K_V_1 channels to include other subunits, especially K_V_1.1 (Brew et al., 2007). Comparability to the human phenotype therefore remains unclear, as the disease-associated K_V_1 reduction are mostly missense variants that can still assemble in heteromers and can exhibit a dominant-negative effect. To address this limitation, we explored the effect of BMK86-P1 in primary cortical neurons. Administration of BMK86-P1 had no effect on passive membrane properties and AP properties. We observed a slight hyperpolarization of the resting membrane potential and a reduction of the AP threshold in both recordings with extracellular solution and BMK86-P1 compared to baseline, suggesting a time-dependent, non-toxin-specific effect. This is consistent with reports showing that dialysis of the intracellular components during prolonged whole-cell recordings can affect neuronal properties over time, underscoring the importance of appropriate control recordings (Komagiri & Kitamura, 2003). We further observed a statistically non-significant, minimal effect on firing frequency, in which BMK86-P1 slightly enhanced the firing rate at lower current injections, whereas at higher current injection it tended to decrease firing. In contrast, unspecific K_V_1 channel blockers have caused excessive firing and spontaneous activity, emphasizing the involvement of K_V_1.2 in dampening neuronal excitability (Bekkers & Delaney, 2001). Bearing in mind, the loss of BMK86-P1-mediated block on K_V_1.2-containing heteromers in heterologously expressed cells, the relatively small impact of BMK86-P1 on neuronal intrinsic excitability and AP characteristics could, in part, be explained by the fact that K_V_1.2 is expressed in complex heteromers with distinct subunit compositions that vary across neuronal types and subcellular compartments (Manganas & Trimmer, 2000; Rasband & Shrager, 2000; H. Wang, Kunkel, Schwartzkroin, & Tempel, 1994). However, incomplete oxidation of BMK86-P1 may reduce the toxińs potency and is therefore to be more likely to explain the unexpectedly minimal effects on neuronal activity.

In fact, we observed minor differences on the network level, where the cumulative distribution of sEPSC amplitude in primary cortical neurons decreased over time during control recordings, whereas amplitudes remained stable at baseline levels during BMK86-P1 treatment. Moreover, we found that the instantaneous sEPSC frequency slightly increased when BMK86-P1 was applied, while remaining unchanged when Extra was applied. It appears that the toxin peptide may partially compensate for time-dependent effects at the neuronal network level. Intriguingly, the role of K_V_1 channels in synaptic transmission has been demonstrated before: The inactivation of K_V_1.2-containing channels at the AIS of layer 5 pyramidal neurons enhance the amplitude of the excitatory postsynaptic potential recorded in postsynaptic cells, while the AP properties, particularly AP width, remained unaffected at the soma of these cells (Kole et al., 2007; Shu, Yu, Yang, & McCormick, 2007). Further, the role of K_V_1 channels in AP repolarization at axon collaterals and presynaptic terminals of neocortical pyramidal cells was underlined, independently of AP properties at the soma (Foust, Yu, Popovic, Zecevic, & McCormick, 2011). Since we recorded at the soma, it would be interesting to ascertain whether BMK86-P1, despite its minimal blocking effect on K_V_1.2 heteromers, has an impact on axonal APs, in a region with high K_V_1.2/K_V_1.1 heteromer expression (Rasband & Shrager, 2000), and if this effect is causative for the slightly increased EPSC amplitude observed. This may open an opportunity for a mild therapeutic effect for patients with GOF variants in *KCNA2*. Nonetheless, the likely incomplete oxidation resulting in reduced toxin affinity may mask the actual effect on network function and is the cause for the subtle change. Here further optimisation in the control of the oxidative folding process of cysteine-rich peptides will be needed in the future.

In summary, we showed that BMK86-P1 blocks selectively K_V_1.2-containing channels and loses its potency when assembled in heteromeric K_V_1.2 channel complexes. However, it is challenging to draw definitive conclusions, as the overall toxin effect is very subtle, potentially due to incomplete oxidation. Importantly, our results support that the toxin affinity depends on amino acid residue p.V381, as this effect can be reversed by introducing the K_V_1.1 homologous substitution p.V381Y, bearing in mind changed activation properties. Thus, our work proposes the potential of BMK86-P1 as a potential pharmacological tool whose K_V_1.2-inhibitory effect can be controlled by a single amino acid in the channel pore. Combining V381Y together with disease-associated K_V_1 variants could allow variants to be investigated in physiologically relevant systems, such as neurons, especially if further research optimizes oxidation of peptides.

## Author Contributions

Conceptualization, PMW, UBSH & HK; toxin synthesis and quality control, HK; CHO electrophysiology, EB, LO, HL, SI, RL & PMW; HEK electrophysiology, EB, RL; data curation, LO, EB & PMW; data analysis, EB & PMW; writing—original draft preparation, EB & PMW; writing—review and editing, UBSH; visualization, EB & PMW. All authors have read and agreed to the published version of the manuscript.

## Acknowledgments

We thank Mahmoud Koko who provided us with the plasmids for KCNA1 and KCNA2. Sketches in Fig. 3 were partially created via https://app.biorender.com/.

## Data Availability Statement

The data that support the findings of this study are available from the corresponding author upon reasonable request. Some data may not be made available because of privacy or ethical restrictions.

