## Supplemental Files for "Scorpion toxin peptide BMK86-P1 achieves mutation-reversible inhibition of KCNA2 at the cost of reduced efficacy in heteromers and murine neurons"

***Supplementary Material***

1 **Supplementary**

Modelling the effect of BMK86P1- on different heteromer compositions

Heteromers of K_V_1.1 and K_V_1.2 should assemble following a poisson distribution given similar transducation rates (Fig S1A). We then wanted to estimate whether the toxin would work on all heteromers with K_V_1.2 and to which degree this would have to be the case. Therefore, we chose four scenarios of toxin peptide heteromer interaction:

First, we modelled the scenario where each subunit would contribute additively to the total current inhibition:

Model 1 (‘equal add. contribution’)$\sum_{k=0}^{4} \left( \frac{4}{k} \right)p^{k}\left( 1-p \right)^{4-k}\left( 1-\frac{k\left( 1-E \right)}{4} \right)=0.834$

where *k* is the amount of K_V_1.2 subunits per channel, *E* indicates the effect in current taken from the homomeric experimental data (0.668) and *p* is the chance of the individual subunit to be included in the channel (50%).

The second model considers the effect given that one subunit of K_V_1.2 alone would be sufficient to exert the full effect on current inhibition:

Model 2 (‘all or nothing’)$\sum_{k=1}^{4} \left( \frac{4}{k} \right)p^{k}\left( 1-p \right)^{4-k}E+\left( \frac{4}{0} \right)p^{4}\left( 1-p \right)^{4-0}=0.689$

The third model considers a scenario in which only opposing K_V_1.2 subunits (Al-Sabi et al., 2010) contribute to the total current inhibition:

Model 3 (‘opposing subunits’):

$$\sum_{k=0}^{1} \left( \frac{4}{k} \right)p^{k}\left( 1-p \right)^{4-k}E+\sum_{k=3}^{4} \left( \frac{4}{k} \right)p^{k}\left( 1-p \right)^{4-k}E+\left( \frac{2}{4} \right)p^{4}\left( 1-p \right)^{4-2}(\frac{2}{3}+ \frac{1}{3}E)=0.85475$$

The fourth model represents the case where only the K_V_1.2 homomer contributes to the current inhibition:

Model 4 (‘only homomer’)$\sum_{k=0}^{3} \left( \frac{4}{k} \right)p^{k}\left( 1-p \right)^{4-k}+\left( \frac{4}{04} \right)p^{4}\left( 1-p \right)^{4-4}=0.979$

The results of these models are summarised in Fig. S1B.

1.1 **Supplementary Figures**

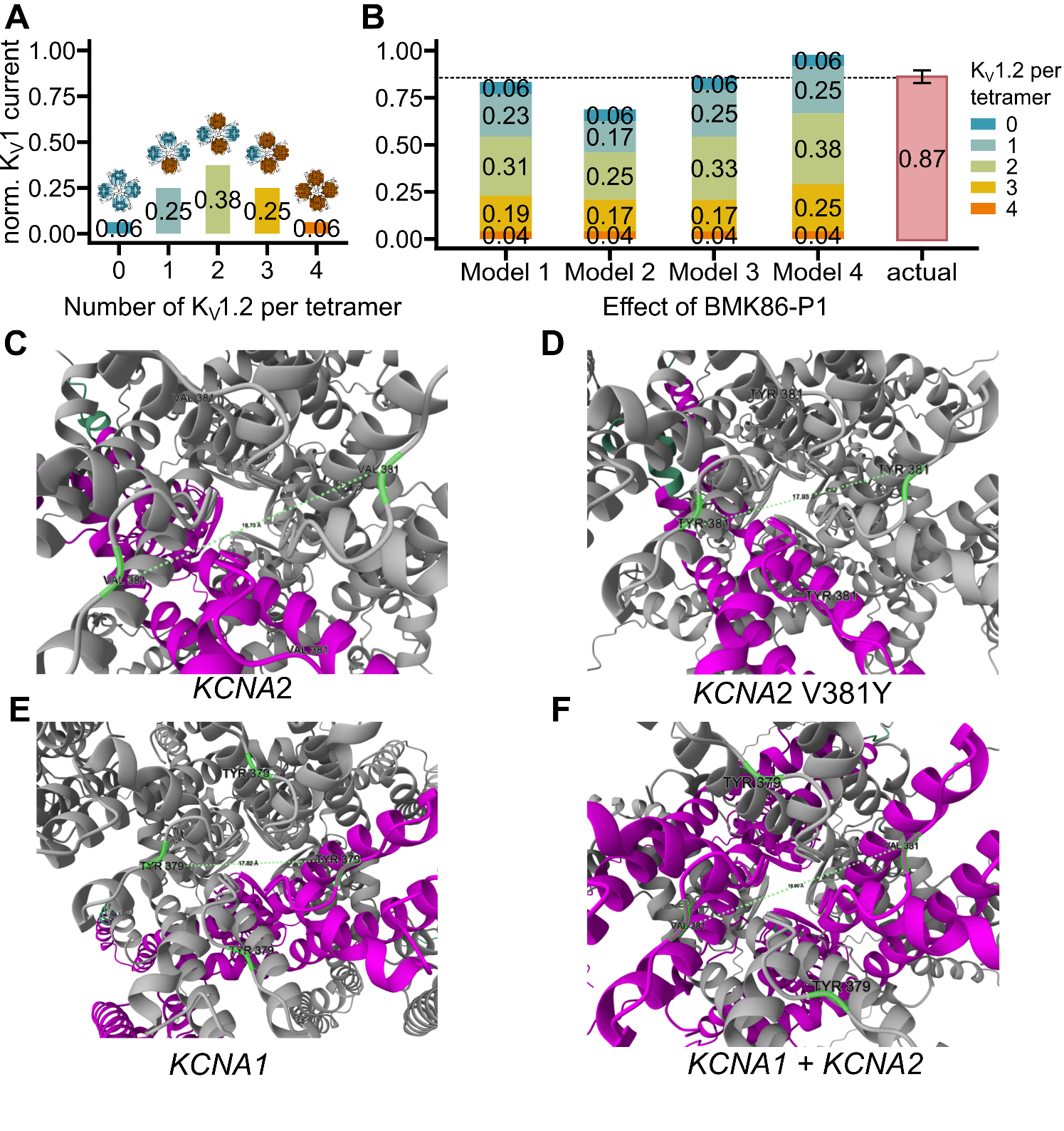

**Supplementary Figure 1** Predicted interaction of BMK86P1 on KV1 heteromers. (A) Theoretical contribution to total K_V_1 current magnitude of different heteromeric compositions. Symbols above indicate the composition with K_V_1.1 in blue and K_V_1.2 in red. (B) Theoretical effect of BMK86-P1 on K_V_1.2/K_V_1.1 current amplitude given different assumptions for the effect of the toxin peptide on heteromers (see text for details on Model 1-4) and recorded effect for *KCNA1*+*KCNA2* (actual).

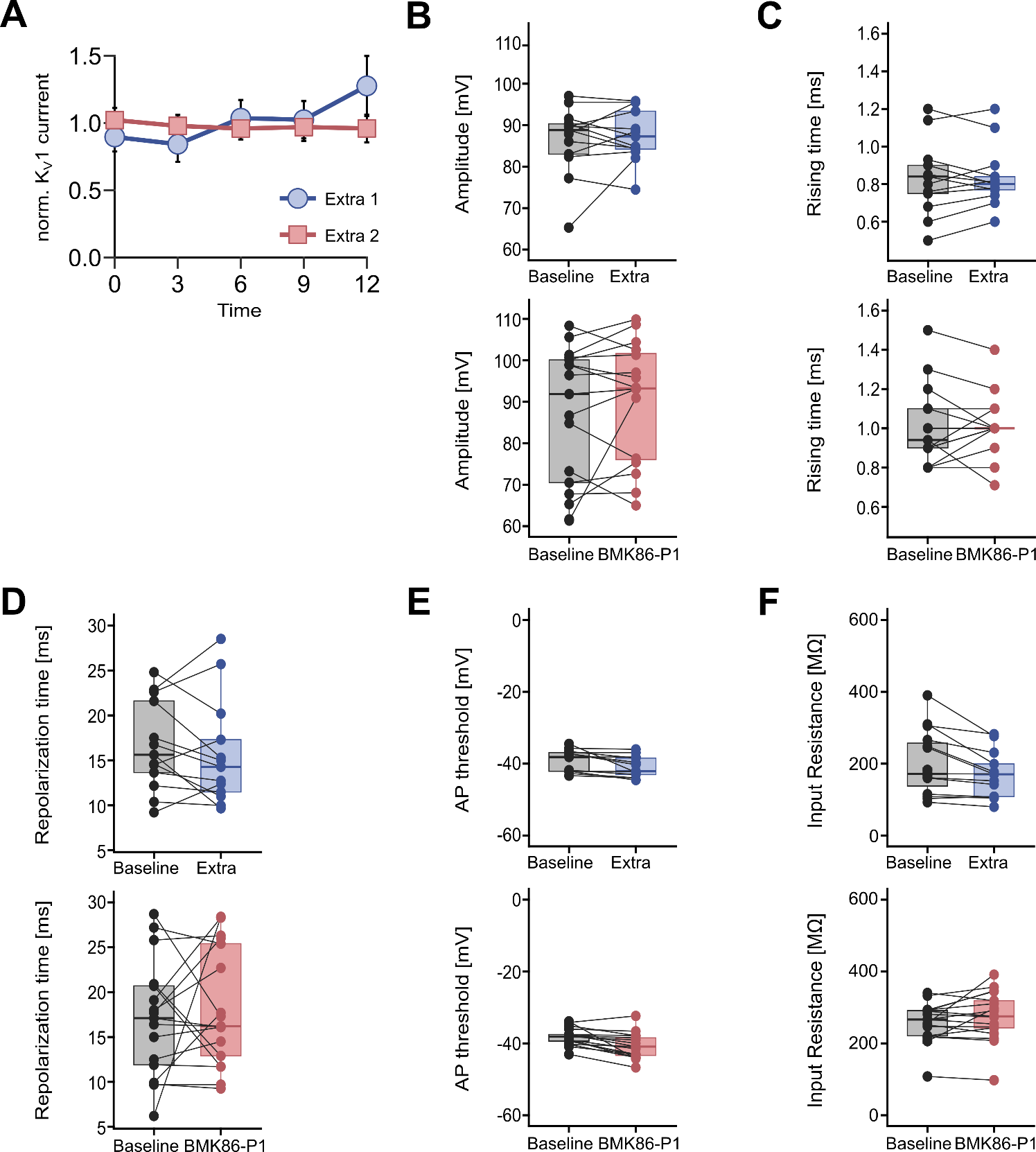

**Supplementary Figure 2.** The effect of BMK86-P1 on AP properties of pyramidal cortical neurons. (A) Normalized potassium currents recorded after two consecutive applications of extracellular solution. (B) Amplitude recorded before (baseline, black) and after application of extracellular solution (blue) or BMK86-P1 (red). (C) Rising time recorded before (baseline, black) and after application of extracellular solution (blue) or BMK86-P1 (red). (D) Repolarization time recorded before (baseline, black) and after application of extracellular solution (blue) or BMK86-P1 (red). (E) AP threshold recorded before (baseline, black) and after application of extracellular solution (blue) or BMK86-P1 (red). (F) Input resistance was compared to baseline (black) following application of extracellular solution (blue) or BMK86-P1 (red). All data are obtained as median, the boxplot shows the interquartile range. All p-values of Wilcoxon-signed rank test were not significant (p > 0.05).
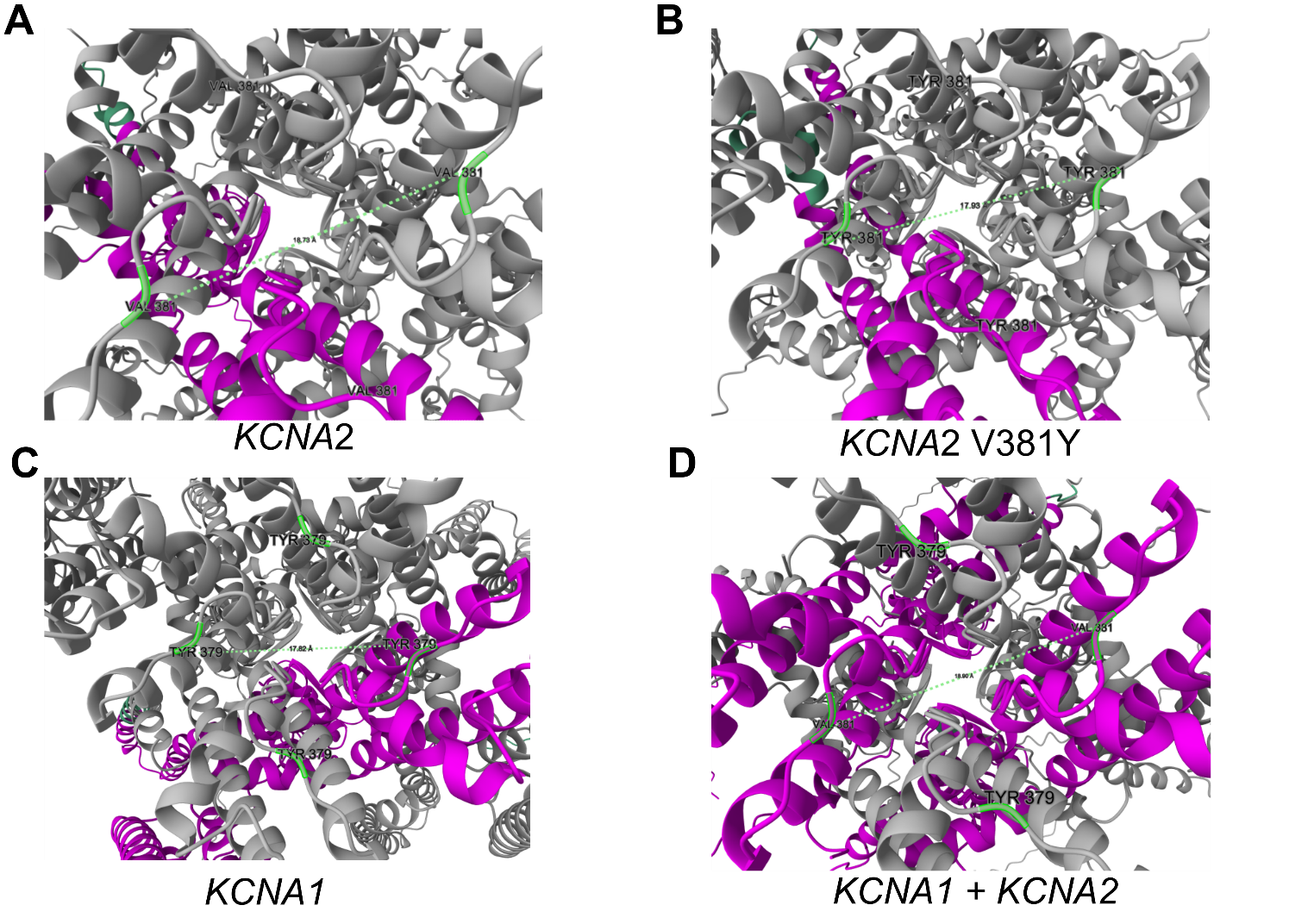

**Supplementary Figure 3**. Alphafold3 models for different KV1 tetramers. (A-D) Zoom in on top of the selectivity filter, pink highlights one KV1 subunit for (B-E) and both KV1.2 subunits for (F). (A) For the homomer of *KCNA2* the distance between opposing p.V381 is 18.73 Å. (B) In the *KCNA2* V381Y tetramer the distance is 17.93 Å. (C) For *KCNA1* the distance between opposing p.Y379 is 17.82 Å. (D) For the heteromer the distance between opposing *KCNA2* p.V381 is 18.90 Å and 17.73 Å for opposing *KCNA1* p.Y379. The plDDT for all labelled amino acids was > 90.
